# Metabolic-scale gene activation screens identify SLCO2B1 as a heme transporter that enhances cellular iron availability

**DOI:** 10.1101/2022.02.06.479330

**Authors:** Gokhan Unlu, Benjamin Prizer, Ranya Erdal, Hsi-Wen Yeh, Erol C. Bayraktar, Kıvanç Birsoy

## Abstract

Iron is the most abundant transition metal in cells and essential for a wide range of biochemical processes. While most mammalian cells take up iron through receptor-mediated endocytosis of transferrin, molecular players involved in iron utilization under iron-limiting conditions are incompletely understood. To address this, we performed several parallel metabolism-focused CRISPRa gain of function screens, which revealed metabolic limitations under stress conditions. Screens for iron restriction identified expected members of iron utilization pathways, but also *SLCO2B1*, a poorly characterized membrane carrier. Expression of *SLCO2B1* is sufficient to increase intracellular iron stores, bypass the essentiality of transferrin receptor-mediated iron uptake and enable cell proliferation under iron restriction. Mechanistically, SLCO2B1 mediates heme-analog import in cellular assays. Heme uptake by SLCO2B1 provides sufficient iron for cell proliferation through heme oxygenases. Notably, *SLCO2B1* is predominantly expressed in microglia in the brain and primary microglia from Slco2b1^−/−^ mice exhibit a strong defect in heme analog import. Altogether, our work identifies SLCO2B1 as a microglia-enriched plasma membrane heme importer and provides a genetic platform to identify metabolic limitations under stress conditions.

## INTRODUCTION

Iron is the most abundant transition metal and essential for almost all living species. A significant fraction (2%) of human proteins binds to iron in the form of heme (iron-protoporphyrin IX), iron-sulfur (Fe-S) clusters, and as free ion (Andreini et al., 2018; Kaplan and Ward, 2013). Owing to its unique redox characteristics, iron mediates a wide range of biological process including oxidative phosphorylation, DNA replication and antioxidant response. In mammals, iron deficiency causes anemia but is also associated with impaired neurological function and weakened immune response (Jiang et al., 2019; Yager and Hartfield, 2002). In contrast, excess cellular iron generates reactive oxygen species that damage cellular macromolecules, and is linked to aging, liver failure and neurodegeneration (Sato et al., 2022; Wang et al., 2017). Most mammalian cells take up iron in the form of transferrin, a serum glycoprotein that maintains it in soluble form and limits the generation of free radicals. However, iron also associates with other carriers in the serum such as ferritin, heme and albumin. These carriers are particularly critical under disease conditions where transferrin-bound iron levels are limiting (Leitner and Connor, 2012; Moos and Morgan, 1998). However, molecular players involved in iron utilization under iron-restricted conditions are incompletely understood.

To address this, we developed a metabolic scale CRISPR activation (CRISPRa) screening platform and performed gain-of-function genetic screens under iron-limiting conditions. These screens revealed multiple mechanisms that enable cell proliferation under iron restriction. Our work also identified *SLCO2B1*, a poorly characterized plasma membrane transporter, as a heme transporter that promotes cellular iron availability. *SLCO2B1* is predominantly expressed in microglia in the brain and primary microglia from *Slco2b1*^−/−^ mice exhibit a strong defect in heme analog import. Together, our results reveal metabolic limitations under iron restriction and identify a plasma membrane heme transporter in microglia.

## RESULTS

### Metabolic-scale gene activation screens identify metabolic processes limiting for cell proliferation under iron restriction

One approach to identify metabolic limitations under stress is to systematically induce the expression of small molecule transporters and enzymes and determine those that restore cell proliferation in response to a particular stress. Activation of target gene expression can be achieved with nuclease dead Cas9 (dCas9a) fused to a transcriptional activator (VP64-p65-Rta). Using this technology, we generated a metabolism focused sgRNA library consisting of 32,509 sgRNAs that target 2989 metabolic genes (**Figure 1A**). To test the robustness of the approach, we first performed gain-of-function genetic screens in dCas9a expressing PaTu-8988t pancreatic cancer cells (**Figure S1A**) treated with near-lethal doses of several metabolic inhibitors. These small molecules include Antimycin A, an electron transport chain (ETC) inhibitor, CB-839, an inhibitor of glutaminolysis, buthionine sulfoximine (BSO), an inhibitor of glutathione synthesis, and palmitate, a saturated fatty acid that disrupts membrane lipid homeostasis at high levels. After selection with these agents, we harvested DNA from surviving cell populations and calculated the percentage of total sgRNA reads mapping to each gene. Genes with highest percentage sgRNA reads protect cells from the corresponding toxic agents and reveal potential metabolic limitations for cell proliferation under these stress conditions.

**Figure 1.**
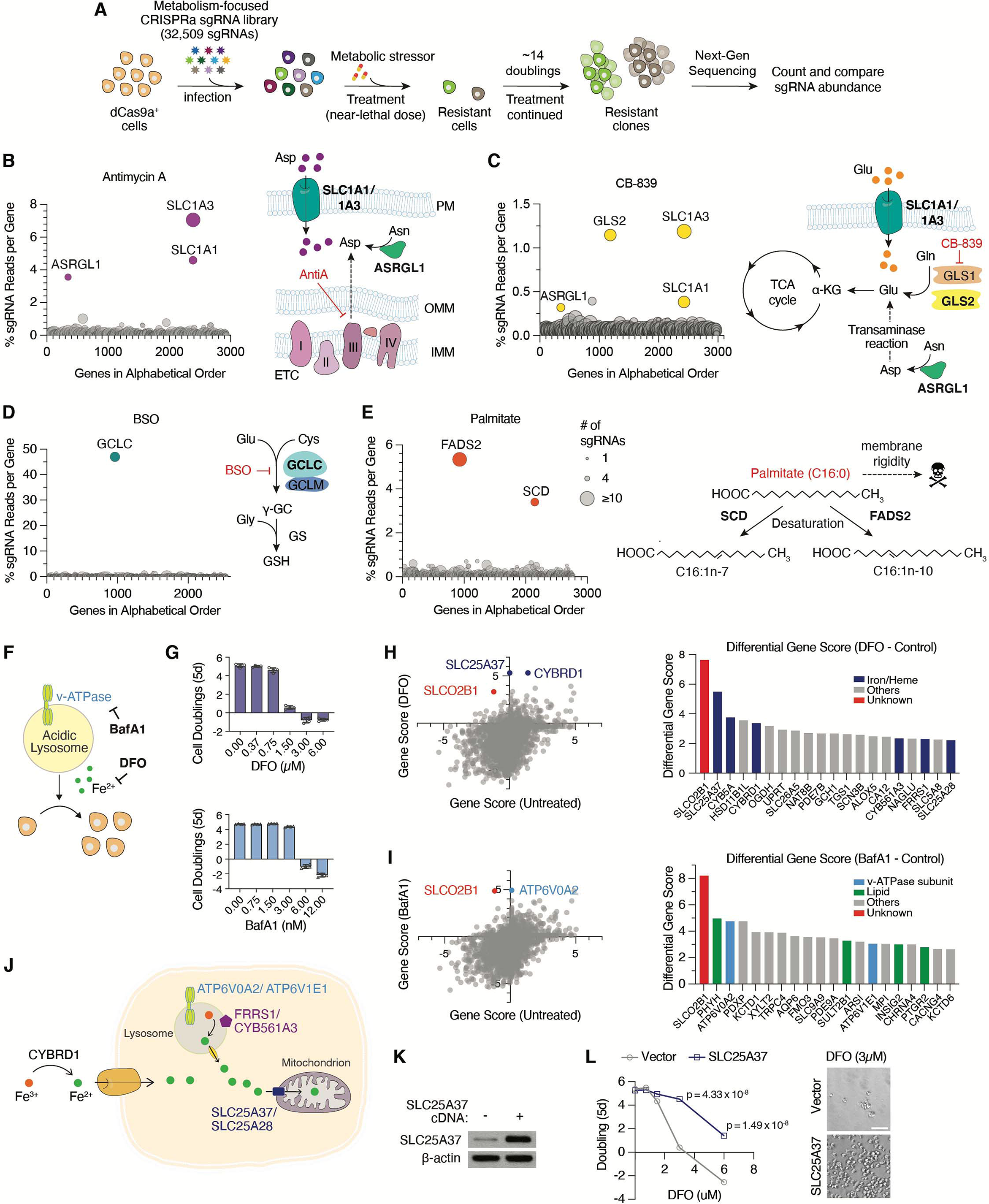
Metabolic scale CRISPRa screens identify processes limiting for cell proliferation under iron restriction. (A) Scheme describing metabolism-focused positive selection CRISPRa screens. (B-E) CRISPRa screen results for (B) Antimycin A (10 nM), (C) CB-839 (6 μM), (D) BSO (2 mM) in Pa-Tu 8988t dCas9a cells and (E) Palmitate (200 μM) in Jurkat dCas9a cells. Data were plotted as genes in alphabetical order vs. percentage of sgRNAs reads per gene. Percent total sgRNA reads was calculated by summing up all sgRNA reads mapped to a gene and calculating the percentage of this sum to the number of sgRNA reads acquired from the entire population. Bubble size indicates number of detected sgRNAs post-screen. (F) Maintaining iron availability is essential for cell proliferation. BafA1 and DFO decreases cellular iron availability through different mechanisms. (G) Dose-dependent effects of DFO and BafA1 on proliferation of Jurkat cells (Mean ± SEM, n = 5). (H) Gene scores of untreated vs. DFO (2 μM)-treated Jurkat-dCas9a cells (left). Gene score is the median log_2_ fold change in the abundance of all sgRNAs targeting that gene during the screening period. Differential gene scores of top 20 genes providing resistance to DFO (right). Iron/heme-related genes are indicated in dark blue. (I) Gene scores of untreated vs. BafA1 (5 nM)-treated Jurkat-dCas9a cells (left). Differential gene scores of top 20 genes providing resistance to BafA1 (right). Vacuolar ATPase subunits are indicated in light blue, lipid metabolism-related genes in green, other genes in gray. (J) Scheme displaying selected scoring genes conferring resistance to iron-restriction. (K) Immunoblot analysis of control and *SLC25A37*-overexpressing Jurkat cells. β-actin was used as loading control. (L) Fold change in the number (log_2_) of control and *SLC25A37*-overexpressing cells after 5 days in the presence of indicated DFO concentrations (left). Representative bright-field micrographs of Jurkat cells after 5-day treatment with 3 μM DFO (right). Scale bar is 50 μM.

Among the scoring genes that protect cells against ETC inhibition are *SLC1A1* and *SLC1A3*, both of which are aspartate/glutamate plasma membrane transporters and increase aspartate import from the culture media. Indeed, previous work has determined that aspartate is a limiting metabolite for cancer cell proliferation under ETC inhibition (Birsoy et al., 2015). Interestingly, our screens also identified *ASRGL1*, a putative human L-asparaginase, which catalyzes the deamidation of asparagine to produce aspartate (**Figure 1B**). While ASRGL1 has previously been shown not to display amidase activity at physiological levels of expression (Pavlova et al., 2018), our data suggest that ASRGL1 may synthesize meaningful levels of aspartate when overexpressed in human cells. Conferring resistance to CB-839, our screens identify *GLS2*, a paralog of *GLS1*, as a gene that can bypass GLS1 inhibition, supporting the selective on-target effect of CB-839 on GLS1 (**Figure 1C**). *SLC1A1/3* and *ASRGL1* also scored likely due to their ability increase the cellular levels of glutamate, a limiting metabolite for TCA cycle progression under glutaminase inhibition. In line with the fact that BSO is a competitive glutamate analog (Meister, 1995), the only scoring gene in the BSO screen was glutamate—cysteine ligase catalytic subunit (*GCLC*), which catalyzes the first enzymatic step in glutathione (GSH) synthesis (**Figure 1D**). Finally, expression of *FADS2* (fatty acid desaturase) and *SCD* (stearoyl-CoA desaturase), the two enzymes involved in fatty acid desaturation, protect cells against lipid saturation stress upon palmitate treatment (**Figure 1E**). Additional hits conferring resistance to palmitate were *FADS1*, which introduces *cis* Δ5 double bond during polyunsaturated fatty acid (PUFA) synthesis (Cho et al., 1999); *ACSL4*, the major enzyme that produces long chain PUFA containing lipids, *ADIPOR2*, which acts as a sensor for membrane fluidity and *SLCO4A1*, an uncharacterized transporter (Devkota et al., 2017; Ruiz et al., 2019) (**Figure S1B**). Notably, *SLCO4A1* has previously been shown to be co-essential with enzymes in fatty acid metabolism, indicating its potential role in membrane lipid homeostasis (Aregger et al., 2020). These results highlight the utility of metabolism-scale gain of function screens to discover potential metabolic limitations under stress conditions.

We next sought to use this platform as a discovery tool to identify cellular pathways that regulate cellular iron availability under iron restriction. While iron chelators are commonly used to deplete cellular iron, we recently showed that disrupting lysosomal acidity also decreases cellular iron levels by hindering transferrin-mediated iron import (Weber et al., 2020). We therefore performed positive selection CRISPRa screens in Jurkat dCas9a cells (**Figure S1A**) and determined genes whose expression would overcome anti-proliferative effects of iron restriction induced by an iron chelator, deferoxamine mesylate (DFO, 2 μM), and a lysosomal pH inhibitor, Bafilomycin A1 (BafA1, 5 nM) (**Figures 1F and 1G**). Scoring genes should identify metabolic limitations under iron depletion and reveal alternative pathways involved in iron acquisition. Consistent with the observation that DFO and BafA1 impair cell proliferation via distinct mechanisms, most of the scoring genes were unique to each condition. Among the scoring genes in DFO screens are *SLC25A37* and *SLC25A28*, mitochondrial iron importers (Shaw et al., 2006). Additionally, plasma membrane and endosomal ferric reductases that reduce Fe^3+^ to Fe^2+^ also scored, including *CYBRD1*, *FRRS1* and *CYB561A3* (**Figure 1H**) (Vargas et al., 2003; Wang et al., 2021). In contrast, genes conferring resistance to BafA1 included v-ATPase subunits *ATP6V0A2* and *ATPV1E1* among the top hits. These subunits are adjacent to the BafA1 binding site on the v-ATPase, and their overexpression likely blocks inhibition of v-ATPase activity by BafA1 (**Figure 1I-J**).

Iron plays essential roles in different compartments in cells. In addition to proteins containing heme or iron-sulfur (Fe-S) clusters, whose syntheses occur in the mitochondrial matrix, there are other key enzymes containing iron, but devoid of Fe-S clusters or heme. For example, di-iron center is present in a group of non-mitochondrial enzymes such as ribonucleotide reductases, oxygenases and fatty acid desaturases (Bollinger Jr et al., 1991; Ryle and Hausinger, 2002; Shen et al., 2020). Given that both mitochondrial iron transporters scored in our genetic screens, we hypothesized that iron availability in mitochondria, but not in the cytosol or other organelles, is the limiting process for cell proliferation under environmental iron restriction. Consistent with this hypothesis, overexpression of mitochondrial iron transporter *SLC25A37* was sufficient to sustain cell proliferation at lethal DFO concentrations (**Figures 1K and 1L**). These data suggest that mitochondrial iron availability is the limiting process for proliferation when cells are deprived of iron. Altogether, our screens identify distinct metabolic players involved in cell survival and proliferation under iron chelators and lysosomal pH inhibition. Strikingly, *SLCO2B1*, a poorly characterized small molecule transporter, was the only common hit upon DFO and BafA1-induced iron restriction (**Figures 1H, 1I, S1C, S1D and S1E**). We therefore focused our attention on it.

### SLCO2B1 expression is sufficient to sustain cell proliferation under iron restriction

SLCO2B1 is a member of organic anion transporter family (König, 2011) and has previously been shown to transport a multitude of drugs and steroid hormone conjugates (Grube et al., 2006; Medwid et al., 2019). However, its precise physiological substrate and relevance to iron metabolism is unknown. To begin to study its function, we generated a Jurkat cell line that expresses *SLCO2B1* cDNA (**Figure S1F**). Consistent with the screen results, cells expressing *SLCO2B1* can proliferate even under lethal doses of DFO and BafA1, in contrast to those expressing a control vector, its closely related paralog *SLCO2A1* or a short *SLCO2B1* isoform (**Figures 2A, 2B, S1G, S1H**). This finding is generalizable to other cell lines, as *SLCO2B1* expression in PaTu-8988t cells protected them from DFO and BafA1-induced toxicity (**Figures S1I and S1J**). Additionally, positive selection CRISPRa screens in a mouse pancreas cancer cell line identified Slco2b1 as the top protective gene against lysosomal pH dysfunction (**Figures S1K, S1L, S1M, S1N and S1O**). SLCO2B1-overexpressing cells displayed similar resistance to deferiprone (DFP), a lipid-soluble iron chelator, v-ATPase inhibitor concanamycin (ConA) and NH^4^Cl, which disrupts lysosomal acidity by acting as a lysosomotropic weak base (Klempner and Styrt, 1983) (**Figures 2C, 2D, 2E**). These findings reveal that *SLCO2B1* expression is sufficient to sustain proliferation of mammalian cells under different modes of iron depletion.

**Figure 2.**
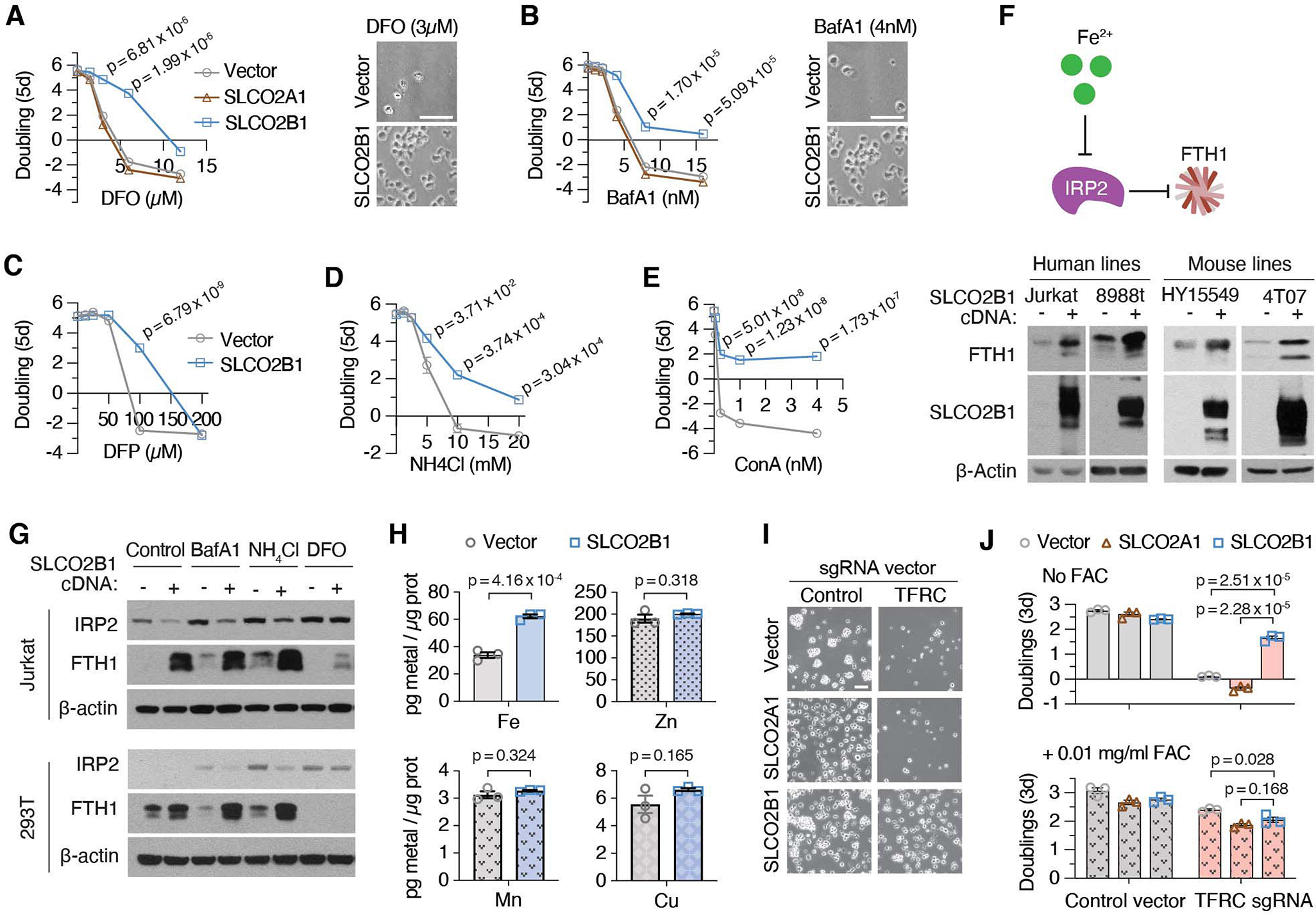
*SLCO2B1* expression is sufficient to promote iron availability and sustain cell proliferation under iron restriction. (A) Fold change in the number (log_2_) of vector, *SLCO2A1* and *SLCO2B1*-expressing cells after 5 days in the presence of indicated DFO concentrations (left). Representative bright-field micrographs of Jurkat cells after 5-day treatment with 3 μM DFO (right). Scale bar is 50 μm. (B) Fold change in the number (log_2_) of vector, *SLCO2A1* and *SLCO2B1*-expressing cells after 5 days in the presence of indicated BafA1 concentrations (left) and representative bright-field micrographs of Jurkat cells after 5-day treatment with 4 nM BafA1 (right). Scale bar is 50 μm. (C-E) Fold change in the number (log_2_) of vector and *SLCO2B1*-expressing cells after 5 days in the presence of indicated (C) DFP, (D) NH^4^Cl and (E) ConA concentrations (F) Schematic (top) showing IRP2-mediated regulation of ferritin heavy chain (FTH1). Increased iron levels inhibit IRP2, a negative regulator of *FTH1* expression. Immunoblot analysis of FTH1 and SLCO2B1 in indicated human and mouse cell lines expressing vector or *SLCO2B1* cDNA (bottom). β-actin was used as loading control. (G) Immunoblot analysis of IRP2 and FTH1 in Jurkat and 293T cells expressing vector or SLCO2B1 cDNA, in the absence or presence of iron depleting agents BafA1 (3 nM in Jurkat; 10 nM in 293T), NH_4_Cl (1 mM in Jurkat; 5 mM in 293T) or DFO (3 μM in Jurkat; 100 μM in 293T). (H) Abundance of indicated metals in Jurkat cells expressing vector or *SLCO2B1* cDNA. Results were normalized to total amount of protein and reported as ‘pg metal per μg protein’. (I) Representative bright-field micrographs of *TFRC* knockout Jurkat cells expressing vector, *SLCO2A1* or *SLCO2B1* cDNA. Scale bar is 50 μm. (J) Fold change in the number (log_2_) of *TFRC* knockout Jurkat cells expressing a control vector, *SLCO2A1* or *SLCO2B1* cDNA without (top) or with 0.01 mg/ml ammonium ferric citrate (FAC) supplementation (bottom) after 3 days. (Mean ± SEM, n= 3, Student’s t-test, 95% confidence interval).

### SLCO2B1 promotes iron availability, independently of transferrin-mediated iron uptake in mammalian cells

We next sought to determine the role of *SLCO2B1* in cellular iron metabolism. To address this in an unbiased way, we profiled cellular metabolites from Jurkat cells expressing *SLCO2A1* (control) or *SLCO2B1* cDNA by liquid chromatography-mass spectrometry (LC-MS). However, only few metabolites change in response to *SLCO2B1* overexpression, suggesting that *SLCO2B1* expression does not cause a strong change in the levels of cellular metabolites (**Figures S2A, S2B and S2C**). Cellular iron homeostasis is tightly controlled in cells by a post-translational mechanism mediated by IRP1/2 pathway (Sviderskiy et al., 2019). An increase in iron levels negatively regulate IRP2, enhancing mRNA stability of ferritin heavy chain 1 (FTH1), the major iron storage protein in mammalian cells. To test whether *SLCO2B1* modulates iron homeostasis, we determined IRP2 and FTH1 protein levels in cells expressing *SLCO2B1*. Expression of *SLCO2B1* caused a marked induction in the protein levels of FTH1 across several mammalian cell lines tested, under standard culture conditions (**Figures 2F and S2D**). We next asked whether *SLCO2B1*-expressing cells maintain higher levels of iron stores even under iron restriction. Indeed, expression of *SLCO2B1* strongly blocked the increase in IRP2 and decrease in FTH1 protein levels in response to treatment with iron chelators or lysosomal pH inhibitors (**Figure 2G**). Finally, to directly measure iron levels in these cells, we conducted inductively coupled plasma mass spectrometry (ICP/MS) measurements. Consistent with the increase in ferritin levels, iron levels in *SLCO2B1*-expressing cells were ~2-fold higher than those of the parental controls. Notably, we did not observe any significant change in the levels of other divalent metals (Cu, Mn and Zn), indicating a specific effect of SLCO2B1 on iron homeostasis (**Figure 2H**). These results suggest that SLC2OB1 regulates the iron response pathway by increasing cellular iron stores.

Under physiological conditions, TFRC-mediated iron uptake is the main route of iron acquisition (Richardson and Ponka, 1997). Indeed, TFRC is a universally essential gene for the proliferation of most cell lines (Tsherniak et al., 2017). To determine whether *SLCO2B1* expression bypasses the requirement of TFRC-mediated iron uptake, we knocked out *TFRC* by CRISPR/Cas9 in Jurkat cells expressing SLCO2B1, its paralog SLCO2A1 or vector control (**Figure S2E**). Consistent with the essential function of TFRC for cell survival, parental Jurkat cells and those expressing SLC2OA1 died upon its depletion. Supplementation of culture media with excess free iron, i.e. ferric iron citrate (FAC), was sufficient to restore cell proliferation. In contrast, *SLCO2B1* expression in Jurkat cells completely restored their survival and proliferation in response to TFRC loss (**Figures 2I and 2J**). These results strongly suggest that *SLCO2B1* promotes iron availability and cell proliferation under iron restriction, independently of transferrin-mediated iron uptake.

### Heme oxygenase is essential for SLCO2B1-induced resistance to iron restriction

To understand how *SLCO2B1* expression enables cell survival under iron restriction, we performed a negative selection CRISPR screen in parental Jurkat cells and those expressing *SLCO2B1*, treated with a normally lethal dose of BafA1. These screens should identify metabolic processes required for *SLCO2B1*-mediated rescue of iron restriction. Additionally, given that *SLCO2B1*-expressing cells tolerate high levels of BafA1 treatment, this provides a unique opportunity to determine metabolic processes that enable cell proliferation under severe lysosomal dysfunction (**Figures 3A, 3B, S3A and S3B**). Indeed, among the scoring genes differentially essential under lysosomal pH inhibition are those involved in cholesterol (*LSS*, *SREBF2*, *ACAT2*, *SQLE*, *FDFT1*) and sphingolipid (*SPTLC1*, *SPTLC2* and *SGMS1*) biosynthesis. Cholesterol and sphingolipids can be synthesized *de novo*, but also acquired from extracellular lipoproteins through lysosomal enzymes. Similarly, cysteine import into lysosomes requires the lysosomal pH gradient. Likely due to the decrease in lysosomal cysteine storage, cells become dependent on cystine uptake mediated by SLC7A11 (**Figures 3C and S3B**) (Pisoni et al., 1990). Further work is needed to determine why several other processes scored as conditionally essential and how they may compensate for lysosomal dysfunction in mammalian cells.

**Figure 3.**
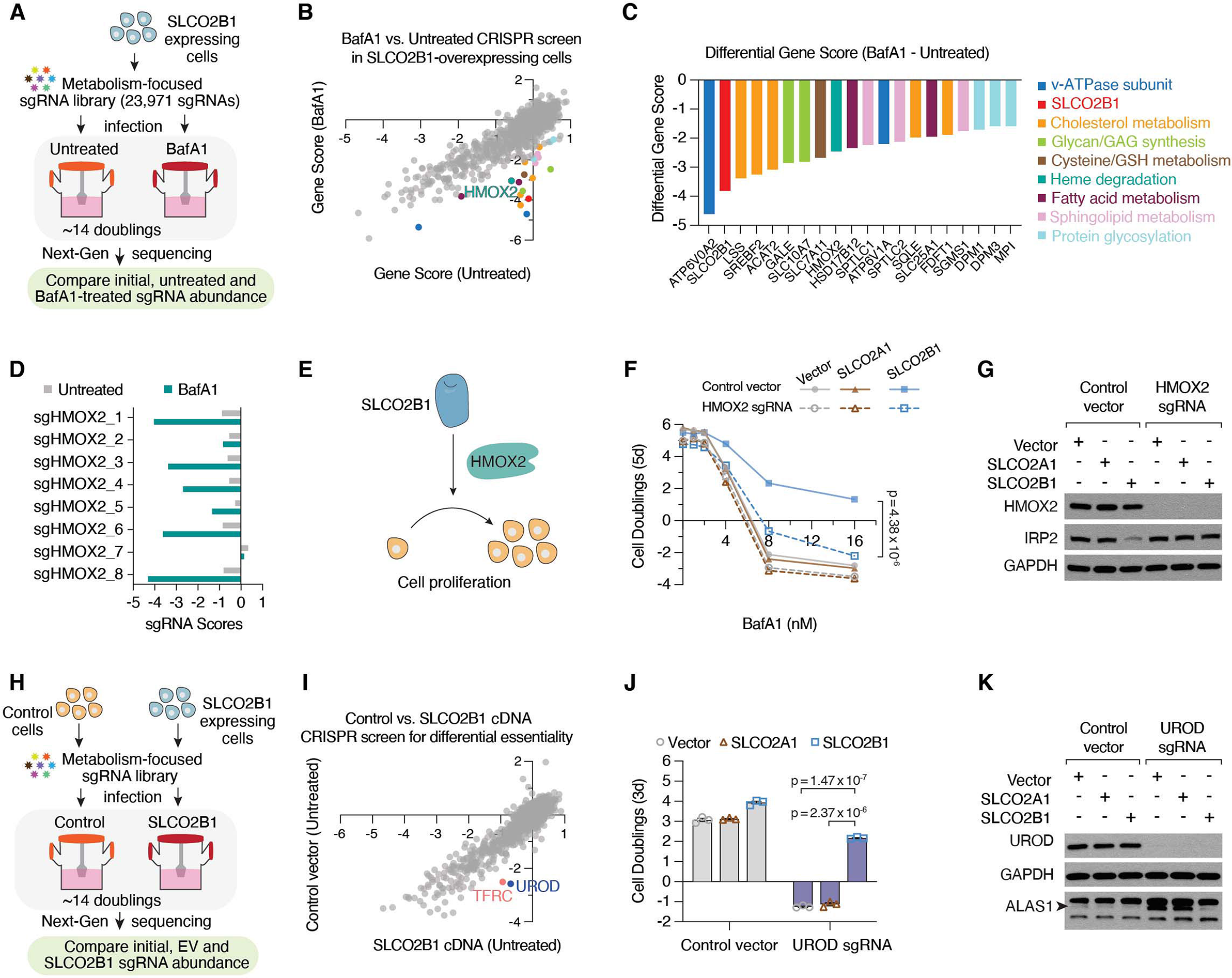
Heme oxygenase is essential for SLCO2B1-mediated resistance to iron restriction. (A) Schematic describing CRISPR loss of function screens in Jurkat cells expressing *SLCO2B1* cDNA in the presence and absence of BafA1. (B) Gene scores of untreated vs. BafA1-treated (4 nM) Jurkat cells expressing *SLCO2B1* cDNA. (C) Top 20 genes scoring as differentially essential upon BafA1 treatment in *SLCO2B1*-expressing cells compared to vector controls. Differential gene scores are plotted, and color matched with metabolic processes they are involved in, as listed on right. (D) sgRNA scores of *HMOX2*-targeting sgRNAs in untreated and BafA1-treated Jurkat cells expressing *SLCO2B1* cDNA. (E) Schematic illustrating the proposed model for SLCO2B1-mediated rescue of cell proliferation under iron restriction. (F) Fold change in the number (log_2_) of control and *HMOX2* knockout Jurkat cells expressing a control vector, *SLCO2A1* or *SLCO2B1* cDNA, during 5-day incubation with or without BafA1 at indicated concentrations. (Mean ± SEM, n= 3, Student’s t-test, 95% confidence interval). (G) Immunoblot analysis for HMOX2 and IRP2 in control and *HMOX2* knockout Jurkat cells expressing vector, *SLCO2A1* or *SLCO2B1* cDNA. GAPDH was used as loading control. (H) Schematic depicting CRISPR loss of function screen in Jurkat cells expressing a vector control compared to those expressing SLCO2B1 cDNA. (I) Gene scores of Jurkat cells expressing a control vector or *SLCO2B1* cDNA. (J) Fold change in the number (log_2_) of control and *UROD* knockout Jurkat cells expressing vector, *SLCO2A1* or *SLCO2B1* cDNA after 3 days. Cells were seeded 7 days after infecting with *UROD* sgRNA vector. (Mean ± SEM, n= 3, Student’s t-test, 95% confidence interval). (K) Immunoblot analysis for UROD and ALAS1 in Jurkat cells expressing vector, *SLCO2A1* or *SLCO2B1* cDNA infected with a control sgRNA or *UROD* sgRNA. GAPDH was used as loading control.

Interestingly, the only scoring gene relevant to iron metabolism and differentially essential in *SLCO2B1* expressing cells was *HMOX2*, a heme oxygenase (HMOX) required for heme degradation (**Figures 3B, 3C, 3D and S3A**). HMOX enzymes break down the porphyrin ring of heme to produce biliverdin, carbon monoxide and iron. Notably, while most mammalian cells express two HMOX paralogs, HMOX2 is the predominantly expressed one in Jurkat cells (TPM of HMOX1 = 0.3, HMOX2 = 32.8, DepMap portal). To test whether *HMOX2* is necessary for *SLCO2B1*-mediated resistance against iron restriction, we deleted *HMOX2* in parental and *SLCO2A1* or *SLCO2B1*-expressing Jurkat cells. Consistent with the screen results, *HMOX2* depletion completely abolished *SLCO2B1*-mediated resistance to BafA1 and DFO treatment (**Figures 3E, 3F and S3C**). Given the requirement of *HMOX2* for *SLCO2B1*-mediated resistance, we reasoned that *HMOX2* loss would block the increase in iron availability and the corresponding decrease in IRP2 protein abundance of *SLCO2B1* expressing cells. In line with our model, *HMOX2* loss restored IRP2 protein levels in SLCO2B1-expressing cells (**Figure 3G**). Collectively, these results suggest that HMOX2 acts downstream of SLCO2B1 to increase intracellular iron availability.

Given that heme degradation is required for SLC2OB1 function, we next sought to determine whether SLCO2B1 is involved in heme metabolism. Interestingly, when we compared gene essentialities between parental cells and those expressing *SLCO2B1*, we found that *UROD* (uroporphyrinogen decarboxylase), like *TFRC*, is essential only in parental cells and not in *SLCO2B1* expressing counterparts (**Figures 3H, 3I, S3E, S3F**). UROD is a universally essential enzyme in mammalian cells and catalyzes the third step of heme biosynthesis in the cytosol. We therefore asked whether *SLCO2B1* expression may bypass the dependence of mammalian cells on *de novo* heme synthesis. While loss of *UROD* blocked the proliferation of parental Jurkat cells, expression of *SLCO2B1* completely eliminated the anti-proliferative effects of *UROD* depletion (**Figure 3J**). Protein levels of ALAS1, the rate limiting enzyme of heme synthesis, is negatively regulated by cellular heme levels. In line with this, expression of *SLCO2B1*, but not control cDNAs, reduced the increase in ALAS1 protein levels in response to UROD depletion (**Figure 3K and S3G**). Similarly, *SLC2OB1* expression blocked the increase in ALAS1 levels in cells treated with succinyl acetone, the pharmacological inhibitor of heme synthesis (**Figures S3D and S3H**). Altogether, these results indicate that *SLCO2B1* expression is sufficient to overcome heme synthesis deficiency.

### SLCO2B1 expression is necessary and sufficient for the uptake of heme analogs

*SLCO2B1*-expressing cells are remarkably resistant to the effects of iron restriction and heme synthesis deficiency, suggesting that SLCO2B1 may be a potential heme importer. To first determine the localization of SLCO2B1, we expressed in PaTu-8988t cells a C-terminally eGFP-tagged version of SLCO2B1, which is fully functional (**Figures S4A, S4B, S4C and S4D**). Confocal microscopy analysis of the SLCO2B1-eGFP fusion protein showed a strong plasma membrane localization (**Figure S4A**). To quantitatively measure steady state heme uptake, we used Zinc mesoporphyrin (ZnMP), a fluorescent heme analog (**Figure 4A**). We treated cells with ZnMP and measured intracellular fluorescence by flow cytometry. ZnMP uptake was markedly higher in SLCO2B1-expressing cells across three human cell lines tested (Jurkat 24h uptake; HepG2 and PaTu-8988t 15 min uptake assays) compared to parental controls (**Figures 4B, 4C and 4D**). To formally prove the direct cellular uptake, we next performed a ZnMP uptake assay in a time- and dose-dependent manner. During a 15-min dose-dependent uptake assay, *Slco2b1*-expressing cells took up significantly more ZnMP at all doses tested (e.g. >5 fold at 0.5 μM) (**Figures 4E, 4F and S4E**). Finally, since heme is a highly redox-active molecule, excess uptake of heme should be more toxic to cells that import it more efficiently. We therefore assayed sensitivity of SLCO2B1-expression and control cells to Hemin, a heme-like porphyrin structure with a ferric chloride center (**Figure 4G**). Indeed, *SLCO2B1* expressing Jurkat cells displayed slower proliferation rates than control and *SLCO2A1*-expressing cells, in response to hemin treatment (**Figure 4H**). Remarkably, upon hemin addition, *SLCO2B1*-expressing cell pellets, in contrast to controls, displayed a discernably brown color, characteristic of hemin accumulation (**Figure 4I**). In parallel, hemin treatment resulted in greater reduction of ALAS1 levels in *SLCO2B1*-expressing cells, suggesting higher hemin availability and subsequent ALAS1 downregulation (**Figure 4J**). These results strongly suggest that SLCO2B1 mediates the transport of heme analogs.

**Figure 4.**
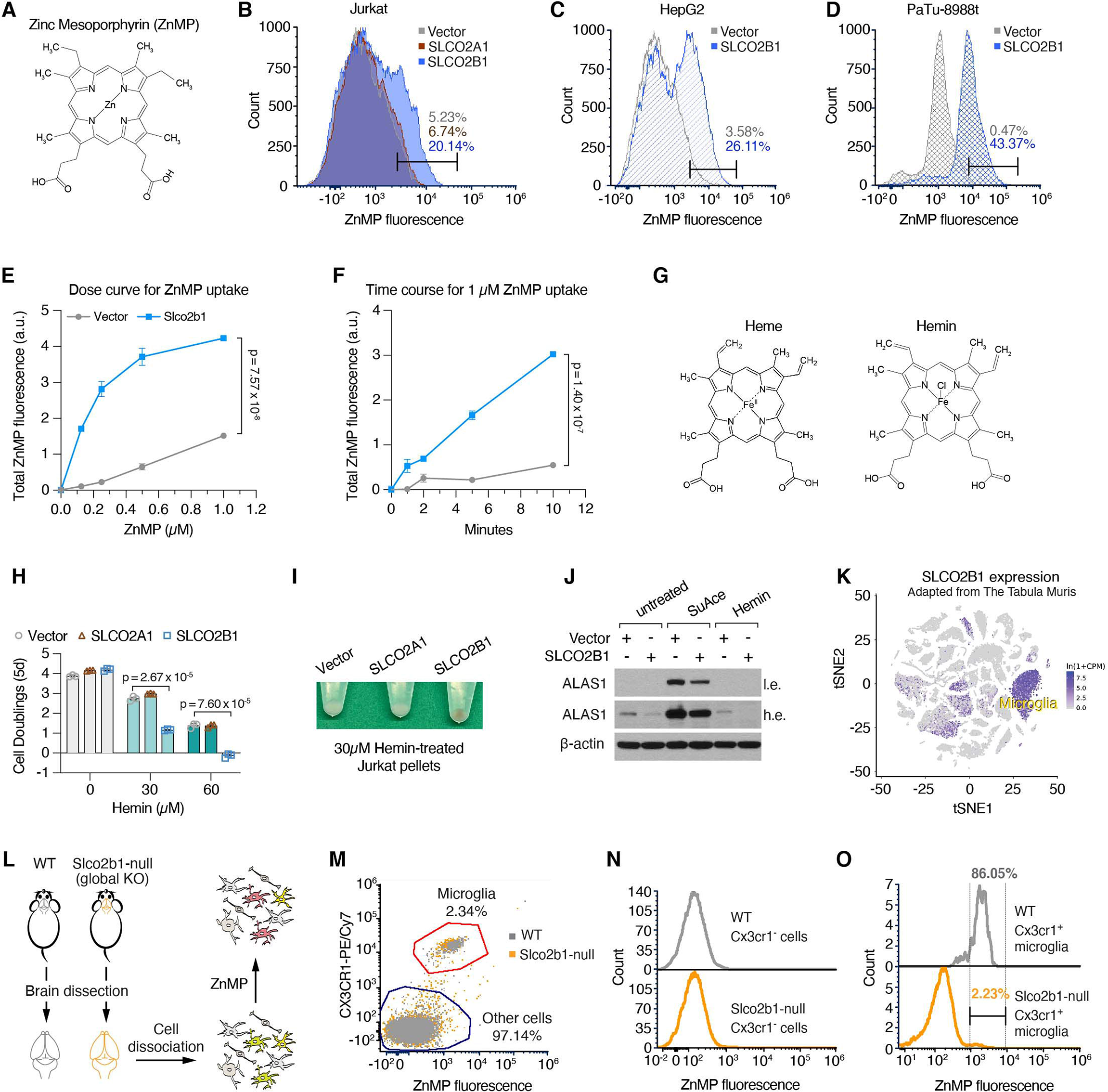
*SLCO2B1* expression is necessary and sufficient for the uptake of heme analogs. (A) Chemical structure of Zinc Mesoporphyrin (ZnMP). (B-D) Flow cytometry analysis of ZnMP uptake in (B) Jurkat cells expressing vector, *SLCO2A1* or *SLCO2B1* cDNA, treated with 5 μM ZnMP for 24 h (C) HepG2 cells expressing vector or *SLCO2B1* cDNA treated with 1 μM ZnMP for 15 min (D) PaTu-8988t cells expressing vector or *SLCO2B1* cDNA treated with 2 μM ZnMP for 15 min. (E) Dose curve for ZnMP uptake in mouse HY15549 pancreas cells expressing vector or *Slco2b1* cDNA treated with indicated doses of ZnMP for 15 min at room temperature. (Mean ± SEM, n= 3, Student’s t-test, 95% confidence interval). (F) Time course for ZnMP uptake in mouse HY15549 pancreas cells expressing vector or *Slco2b1* cDNA treated with 1 μM ZnMP for the indicated time points. (G) Chemical structures of Heme B and Hemin. (H) Fold change in the number (log_2_) of Jurkat cells expressing vector, *SLCO2A1* or *SLCO2B1* cDNA, during 5-day incubation with or without Hemin treatment at indicated concentrations. (Mean ± SEM, n= 3, Student’s t-test, 95% confidence interval). (I) Representative images of cell pellets from Hemin-treated Jurkat cells expressing vector, *SLCO2A1* or *SLCO2B1* cDNA. Cells were treated with 30 μM Hemin for 24 h prior to pelleting. (J) Immunoblot analysis for ALAS1 in Succinyl Acetone (1mM) or Hemin (30 μM) treated Jurkat cells expressing vector or *SLCO2B1* cDNA. β-actin was used as loading control. (K) t-SNE plot for *Slco2b1* expression across single cell transcriptome of mouse cell types. Adapted from Tabula Muris. (L) Experimental strategy for the ZnMP uptake assay of primary cells derived from brains of wild type and *Slco2b1* knockout mice. (M) Flow cytometry analysis of ZnMP fluorescence and Cx3cr1-labeling (microglia marker) in cells isolated from wild-type and *Slco2b1* knockout mouse brains. (N) Flow cytometry analysis of ZnMP fluorescence in Cx3cr1^−^ non-microglial cells (O) Flow cytometry analysis of ZnMP fluorescence in Cx3cr1^+^ microglia

We next sought to test whether *SLCO2B1* is necessary for heme analog import. Notably, most cultured mammalian cells we have tested did not exhibit appreciable levels of SLCO2B1 expression. We therefore analyzed a publicly available single cell transcriptome dataset (Tabula Muris Consortium, 2018) to identify cell types with highest *Slco2b1* expression (**Figure 4K**). Strikingly, *Slco2b1* expression displays a highly restricted expression pattern in microglial cells. To study loss of function of *Slco2b1*, we therefore generated an *Slco2b1*-null mouse model (**Figure S4F**). These knockout mice are viable and enabled us to test the necessity of *Slco2b1* for heme analog uptake in isolated microglial cells (Ledo et al., 2020) (**Figure 4L**). Cells dissociated from brain, other than microglia, did not display detectable ZnMP positivity (**Figures 4M and 4N**), suggesting that microglial cells could indeed be the predominant cell type with heme uptake activity in the brain. Remarkably, this ZnMP uptake activity in wild type microglia almost exclusively depends on *Slco2b1* expression. Indeed, microglia from *Slco2b1* knockout mice display a strong reduction in ZnMP import compared to those from wild type controls (i.e. 86% of wild type microglia, as opposed to 2.23% Slco2b1-null microglia, were ZnMP+) (**Figure 4O**). These results suggest that *Slco2b1* is necessary for heme analog uptake in microglia.

## DISCUSSION

In this study, using a combination of genetic and biochemical strategies, we identify SLCO2B1 as a plasma membrane transporter that mediates heme import. SLCO2B1-mediated heme uptake bypasses two essential metabolic processes in mammalian cells: TFRC-mediated iron uptake and heme biosynthesis. Heme uptake by the heme-responsive paralogs, *hrg-1* and *hrg-4,* has extensively been studied in the context of heme-auxotrophic *C. elegans*. Mammalian homolog of *hrg-1* facilitates lysosomal export of heme in splenic macrophages (Rajagopal et al., 2008; White et al., 2013) and *hrg-4* in worms lacks a mammalian homolog. Furthermore, while previous work proposed FLVCR2 as a plasma membrane heme transporter, its ectopic expression failed to rescue heme-deficiency (Chambers et al., 2021; Duffy et al., 2010). Given that these transporters did not score in our genetic screens, SLCO2B1 may have the highest affinity for heme import among other putative heme importers. Notably, single cell transcriptomics data revealed that *SLCO2B1* expression is highly enriched in microglia (Cao et al., 2020; Tabula Muris Consortium, 2018), indicating that SLCO2B1 might mediate microglial heme import. As free heme levels increase in disease conditions like hematoma and intracranial hemorrhage (Mracsko and Veltkamp, 2014; Vasconcellos et al., 2021), microglial heme uptake by SLCO2B1 could be a defense mechanism to protect neurons and astrocytes from oxidative heme damage. Furthermore, heme uptake may also be relevant in the context of neurodegenerative diseases, as induced *HMOX1* expression and iron accumulation in microglial cells have been observed in Alzheimer’s and Parkinson’s disease (Fernández-Mendívil et al., 2021; Kenkhuis et al., 2021). Future work is needed to determine the precise role of microglial heme uptake in disease conditions.

## EXPERIMENTAL MODEL AND SUBJECT DETAILS

### Mice

All animal studies were performed according to a protocol approved by the Institutional Animal Care and Use Committee (IACUC) at Rockefeller University. Animals were housed in ventilated caging on a standard light-dark cycle with food and water *ad libitum*. The mouse strain C57BL/6N-Slco2b1^tm1a(KOMP)Wtsi^/Mmucd (MMRRC line: 049765-UCD) was recovered from cryopreserved sperm obtained from Mutant Mouse Resource and Research Center (MMRRC) at University of California, Davis. C57BL/6N-Slco2b1^tm1a(KOMP)Wtsi^/Mmucd sperm was crossed to C57BL/6J wild-type strain (from The Jackson Laboratory) through in vitro fertilization at Rockefeller University Transgenic and Reproductive Technology Center. The progeny is maintained under standard conditions and identified through PCR amplification-based genotyping method provided by MMRRC. Homozygous mutant and homozygous wild type animals of 5-8 weeks were used in microglia isolation experiments. Genotypes were revealed to blinded investigators post-analysis.

### Cell Lines, Compounds and Constructs

Jurkat, HepG2, HEK293T and MDA-MB-231 cell lines were purchased from ATCC. Other cell lines were kindly provided by the following investigators: PaTu-8988t pancreatic adenocarcinoma cell line by Dr. Monther Abu-Remaileh (Stanford University, CA), the HY15549 mouse pancreatic cancer line by Nabeel El-Bardeesy (Massachusetts General Hospital Cancer Center, MA), AK196 mouse pancreatic cancer line by Haoqiang Yang (MD Anderson Cancer Center, TX), 4T1 and 4T07 mouse breast cancer cell lines by Dr. Sohail Tavazoie (Rockefeller University, NY). All cell lines were verified to be mycoplasma contamination-free and authenticated by STR profiling. Reagents, compounds, and antibodies used in this study are listed in Table 1.

### Cell Culture Conditions

Unless otherwise indicated, all listed cell lines, except for HepG2, were cultured in RPMI-1640 media (GIBCO) containing 2mM glutamine, 10% fetal bovine serum (SAFC, Sigma Aldrich) and 1% penicillin and streptomycin (Invitrogen). HepG2 cells were cultured in DMEM (GIBCO) containing 4.5g/L glucose, 110mg/L pyruvate, 4mM glutamine, supplemented with 10% fetal bovine serum and 1% penicillin and streptomycin. All cells were maintained at 37°C, 21% O^2^ and 5% CO_2_. For Figure S3, Jurkat cells were cultured in RPMI1640 supplemented with dialyzed FBS (GIBCO #26400-044).

## METHOD DETAILS

### Generation of overexpression and knockout constructs

Gene fragments for coding sequences of human *SLCA25A37*, *SLCO2B1*, *SLCO2A1*; and mouse *Slco2b1* were purchased from Twist Biosciences, then cloned into pLV-EF1a-IRES-Blast vector (Addgene #85133) by Gibson assembly method. pLV-EF1a-IRES-Blast was used as a control vector control in overexpression experiments. Knockout cells were generated with CRISPR/Cas9 method. Forward and reverse oligos targeting *TFRC*, *UROD* and *HMOX2* were annealed and ligated into BsmBI-linearized pLentiCRISPR v2 vector.

### Cell proliferation assays

2,000 Jurkat cells or 500 PaTu-8988t or 500 mouse KPC cells/ per well were seeded, in triplicates, in 0.2 mL RPMI-1640 medium containing indicated treatments in 96-well plates.

On the day of seeding and the final day of treatment (as indicated in corresponding figures), 40 μL of CellTiter-Glo reagent (Promega) was added, then, luminescence was measured on a SpectraMax M3 plate reader (Molecular Devices). Data are presented as cell doublings or the log2 fold change in luminescence on final treatment day compared to initial reading on the day of seeding.

### Generation of knockout and overexpression cell lines

For generation of knockout cells, VSV-G and Delta-VPR lentiviral packaging vectors were simultaneously transfected into HEK293T cells along with plentiCRISPR v2 vector expressing Cas9 and the gene-targeting sgRNA, using XtremeGene9 transfection reagent (Roche). Similarly, for overexpression, pLV-EF1a-IRES-Blast vector containing the gene-of-interest was transfected along with lentiviral packaging vectors VSV-G and Delta-VPR. 60 h post-transfection, the supernatant was collected after passing through a 0.45 μm syringe filter. For transduction, 1 x 10^5^ cells were plated in 6-well plates containing 4 μg/mL polybrene and virus, and then spin-infected by centrifugation at 2,200 rpm for 80 minutes. Mixed population knockouts were selected with puromycin; overexpression cells were selected with blasticidin. Knockout and/or overexpression efficiency was assessed via immunoblotting. Jurkat or PaTu-8988t cells expressing dCas9-VPR were generated following a similar lentiviral transduction strategy. Instead, Edit-R Lentiviral CRISPRa dCas9-VPR vector (Horizon Discovery) was used as the overexpression vector. Single cell clones overexpressing dCas9-VPR were generated by FACS sorting single cells on a BD FACSAriaII, 72-96 h post-infection into 96 well plates and grown for 2 weeks. Clones with strong overexpression were identified by Cas9 immunoblotting.

### CRISPR/Cas9 genetic screens

For metabolic scale CRISPRa screens in human cell lines, a metabolism-focused sgRNA library was designed and screens were performed as previously described (Birsoy *et al.*, 2015; Wang et al., 2015). 2,989 metabolic genes were targeted with a total of 32,460 CRISPRa sgRNAs, whose designing approach was described previously (Horlbeck et al., 2016), and 49 non-targeting sgRNAs were included as controls. Oligonucleotides containing sgRNA sequences were synthesized by Agilent Technologies, PCR amplified and cloned into lentiGuide-Puro vector (Addgene #52963). Briefly, amplicons were inserted into BsmBI-linearized lentiGuide-Puro vector by Gibson Assembly (NEB). Then, Gibson Assembly products were transformed into *E. coli* 10G SUPREME electrocompetent cells (Lucigen). This plasmid pool was used to produce lentivirus-containing supernatants in HEK293T cells. The titer of lentiviral supernatants was determined by infecting target cells at several amounts of virus in the presence of polybrene (4 μg/mL), counting the number of puromycin-resistant infected cells 3 days post-selection. For CRISPRa positive selection screens, 4 million target cells were infected at an MOI of ~0.7 and selected with puromycin (4 μg/mL) 72 h post-infection. An initial pool of 4 million cells was harvested for genomic DNA extraction. The remaining cells were cultured for 14 doublings under specified drug treatment or untreated control conditions. On the final day of screening, cells were harvested for genomic DNA extraction. sgRNA inserts were PCR amplified, purified, and sequenced on MiSeq platform (Illumina). Sequencing reads were aligned to the library of sgRNA sequences and the abundance of each sgRNA was tallied. Results were reported as percentage of total sgRNA reads acquired per gene, gene score or differential gene score. Percent total sgRNA reads was calculated by summing up all sgRNA reads mapped to a gene and calculating the percentage of this sum to the number of sgRNA reads acquired from the entire population. sgRNA scores, representing the log2 fold change of the normalized final read count of the sgRNA from the initial read count of the sgRNA, were calculated. Gene score is defined as the median log_2_ fold change of all sgRNAs targeting the gene in the abundance of all sgRNAs targeting that gene, between the initial and final population. The differential gene score refers to the difference between the gene scores of drug-treated and untreated control groups.

For metabolic scale CRISPRa screens in mouse AK196 pancreas cell line, a metabolism-focused sgRNA library was designed and screens were performed similarly, as described above. For these screens, 1,839 metabolic genes were targeted with a total of 17,031 CRISPRa sgRNAs.

Metabolism-focused CRISPR knockout screens in human lines were performed as described previously (Zhu et al., 2019). 3 x 10^7^ cells were infected with sgRNA library containing 23,971 sgRNAs targeting metabolic genes. Abundance of sgRNAs was quantified post-sequencing on NextSeq500 platform (Illumina). Full results of all CRISPR screens can be found in the supplemental tables S1-S15.

### Metabolite Profiling

For bulk metabolite profiling, 1 x10^6^ Jurkat cells, in triplicate, were grown for 24 h under DFO-, BafA1-treatment or untreated conditions. Then, cells were rinsed in 0.9 % NaCl twice. Polar metabolites were extracted in 80 % methanol containing ^15^N and ^13^C fully-labeled amino acid standards (MSK-A2-1.2, Cambridge Isotope Laboratories, Inc). Extracts were shaken for 10 min with a vortexer, spun at 19,000 g to remove insoluble cell debris, nitrogen-dried and stored at −80°C until liquid chromatography-mass spectrometry analysis (LC-MS). Then, LC-MS was performed as previously described (Garcia-Bermudez et al., 2018). Relative metabolite abundances were quantified using XCalibur QualBrowser 2.2 and Skyline Targeted Mass Spec Environment (MacCoss Lab) using a 5 ppm mass tolerance and a pooled-library of metabolite standards to verify metabolite identity. Relative metabolite levels were calculated by normalizing to total protein levels, as measured by Bicinchoninic Acid Assay (BCA).

### Measurement of Metals by Inductively Couple Plasma / Mass Spectrometry (ICP/MS)

3 x10^7^ Jurkat cells were seeded in triplicates and grown in standard culture conditions for 24 h. Cell pellets were collected and stored at −80°C until ICP/MS analysis. The cell pellet was resuspended in pure, distilled water of ≥80 μl. The cell pellet was lysed on ice, using 5 pulses with a cell disruptor/sonicator microtip. For each ICP/MS replicate, 30 to 50 μl of cell lysate was digested using a 5:1 mixture of nitric acid (OPTIMA grade, 70%, Fisher Scientific) and ultrapure hydrogen peroxide (ULTREX II, 30%, Fisher Scientific). 30 lysate, 500 uL of nitric acid and 100 uL hydrogen peroxide are added to 2 mL polypropylene tubes. This mixture was allowed to digest overnight at room temperature, heated at 95°C just until dry, and resuspended overnight in 1 ml of 2% nitric acid for analysis. The mixture was mixed well the following day to ensure resuspension, and 800 μl more 2% nitric acid is added along with 200 μl of 10X internal standard (prepared in 2% nitric acid). For ICP/MS run, an Agilent 7900 ICP/MS instrument was operated in helium (He) collision cell gas mode for all measurements. Elements were measured at the following isotopes: 56Fe, 55Mn, 63Cu, 66Zn. Calibration standards and samples were prepared in an acid matrix of 2% OPTIMA Grade Nitric Acid. Solutions of Agilent Multi-element Calibration Standard 2A were prepared to obtain an eight-point calibration curve. Agilent Germanium (or Scandium) Standard(s) were added to calibration standards, blanks, and samples. Standards were used to correct for potential sample matrix and/or nebulization effects.

Amounts of metals within a sample were normalized by total amount of protein, as determined by BCA assay (Thermo Fisher Scientific).

### Immunoblotting

1 x 10^6^ cells were washed in cold PBS and lysed in a buffer containing 10 mM Tris-HCl pH 7.4, 150 NaCl, 1 mM EDTA, 1% Triton X-100, 2% SDS, 0.1% CHAPS, and protease inhibitors (Milipore Sigma). Lysates were sonicated, centrifuged at 1,000 g, and supernatant was collcted as the protein lysate. Total protein quantified using BCA Protein Assay Kit (Thermo Fisher) with bovine serum albumin as a protein standard. Protein samples were resolved on 8% or 10%–20% SDS-PAGE gels (Novex, ThermoFisher) and analyzed by standard immunoblotting protocol. Briefly, PVDF membranes (Milipore) were incubated with primary antibodies at 4°C overnight. After washing off the primary antibodies in tris buffered saline/ 0.1% Tween-20 (TBS-T), secondary antibody incubation was performed at room temperature for 1h. Secondary antibodies including anti-mouse IgG–HRP linked (Cell Signaling, 7076) and anti-rabbit IgG–HRP linked (Cell Signaling, 7074), were used at 1:3,000 dilution. Blots were developed by ECL Chemiluminescent detection system (Perkin Elmer LLC) and film exposure. SRX-101A Film Processor (Konica Minolta) and Premium autoradiography Films (Thomas Scientific) were used for developing.

### Immunocytochemistry and Confocal Microscopy

PaTu-8988t cells were seeded on coverslips and grown overnight at 37°C with 5% CO_2_. Next day, cells were fixed in 4 % PFA at room temperature for 15 min, permeabilized in 1% triton X-100 in PBA, and then, blocked in 1% bovine serum albumin (Sigma) for 30 min at room temperature. Primary antibody incubation was performed overnight at 4°C. Then, cells were rinsed in PBS twice and secondary antibody incubation was performed at room temperature for 30 min. After washing twice in PBS, DAPI, nuclear counterstain, incubation was performed at room temperature for 10 min, then cells were washed twice in PBS and mounted in ProLong Gold antifade mountant (Molecular Probes). Slides were imaged with Nikon A1R MP multiphoton microscope with confocal modality, using Nikon Plan Apo γ 60X/1.40 oil immersion objective.

### Real-time quantitative PCR

Total RNA was isolated from mouse livers using TRIzol reagent (Thermo Fisher Scientific), following manufacturer’s manual. After DNase I treatment (New England Biolabs), 1 μg total RNA was used for cDNA synthesis with Superscript III RT kit (Invitrogen). qPCR was performed on a Thermo QuantStudio 6 Flex Real-Time PCR machine. The primer sequences were listed in Table 1. Gene expression levels were normalized to beta-actin using ΔΔCt method.

### Zinc Mesoporphyrin (ZnMP) Uptake and Flow Cytometry Analysis

Adherent cells (HepG2, PaTu-8988t and KP pancreas) were trypsinized and washed with 1X Hank’s Balanced Salt Solution (HBSS) (Milipore Sigma) twice. 1 x10^5^ cells were placed in a microfuge tube and incubated with the indicated concentration of ZnMP (Frontier Scientific) in 1x HBSS for 15 min at room temperature. Uptake was terminated by placing cells on ice. Then, cell pellets were collected, washed twice in 1X HBSS and resuspended in cold FACS buffer (DPBS + 2% FBS + 5mM EDTA). ZnMP uptake was assessed by Flow Cytometry (Attune NxT Flow Cytometer, Thermo Fisher). 10,000 events were recorded per each sample. ZnMP signal was detected by exciting with 561 nm laser, AlexaFluor 568 emission track was recorded as it overlaps with ZnMP emission. Data were analyzed and plotted with FCS Express 7 Research software, version 7.12 (De Novo Software, Inc).

For ZnMP uptake in Jurkat cells, 5μM ZnMP was diluted directly in culture media (RPMI 1640 + 10% FBS). Cells were treated with ZnMP overnight at standard culture conditions (37°C, 5% CO_2_). After that, Jurkat cells were washed twice in PBS and resuspended in FACS buffer. Flow cytometry and data analyses were carried out as described above.

For dose and time-dependent ZnMP uptake assays, mouse KP pancreas cells were used. Cells, in triplicate, were treated with 1:2 serial dilutions of ZnMP, between 0.125 μM and 1μM, including 0 μM control (equal volume DMSO), for 15 minutes at room temperature. For time course uptake assay, cells, in triplicate, were treated with 1 μM ZnMP for 0, 1, 2, 5 and 10 minutes at room temperature. Median fluorescence value of ZnMP^+^-gated population and number of gated events were multiplied to calculate total ZnMP fluorescence. Results were plotted in Prism 9 (GraphPad) as dose or time vs. total ZnMP fluorescence.

### Mouse microglia isolation and ZnMP uptake assay

Mouse microglial cells were isolated from 8-10 week-old female mice as previously described (Ledo *et al.*, 2020). Briefly, mice were sacrificed, whole brains were removed and placed in DPBS (Ca^2+^ and Mg^2+^ free) containing 5% FBS and 1mM HEPES pH7.4. Brain tissue was minced with scissors and incubated in 4,000 U/ml of collagenase D (Roche) at 37°C for 30 minutes. Collagenase digestion was stopped by adding 10 mM EDTA and incubating for additional 5 minutes at 37°C. Digested tissue was passed through 70-μm cell strainer, centrifuged at 2,000 rpm, washed in DPBS, and centrifuged in 38% Percoll gradient for 30 min. Cell pellet containing microglia fraction was washed and resuspended in cold FACS buffer (DPBS + 2% FBS + 5mM EDTA). Non-specific binding was blocked by incubation with Fc blocking antibody (BD Biosciences, Clone 2.4G2) for 15 min. Cells were washed in FACS buffer and stained with Cx3cr1 antibody (Clone: SA011F11) conjugated to PE/Cy7. After that, cells were washed twice in FACS buffer to remove unbound, excess antibody. 5 x10^5^ cells were placed in a microfuge tube and incubated with 2 μM ZnMP (Frontier Scientific) in 1x HBSS for 15 min at room temperature. Uptake was terminated by placing the cells on ice. Then, cell pellets were collected, washed twice in 1X HBSS and resuspended in cold FACS buffer (DPBS + 2% FBS + 5mM EDTA).

ZnMP uptake was assessed by Flow Cytometry (Attune NxT Flow Cytometer, Thermo Fisher). 10,000 events were recorded per each sample. ZnMP signal was detected by exciting with 561 nm laser, AlexaFluor 568 emission track was recorded. Cx3cr1-PE/Cy7 signal was detected by exciting with 488 nm laser and recording emission at 780 nm. Data were analyzed and plotted with FCS Express 7 Research software, version 7.12 (De Novo Software, Inc).

## Supporting information

Table 1. Reagents

## ACKNOWLEDGEMENTS

We thank all members of the Birsoy laboratory for helpful suggestions. G.U. is a Damon Runyon Fellow, supported by the Damon Runyon Cancer Research Foundation (DRG-2431-21), and by the grant UL1 TR001866 from the National Center for Advancing Translational Sciences (NCATS, National Institutes of Health (NIH) Clinical and Translational Science Award (CTSA) program. K.B. is supported by the NIH/NCI (DP2 OD024174-01), NIH/NIDDK (R01 DK123323-01), Pershing Square Sohn Foundation and Mark Foundation Emerging Leader Award; and is a Searle and Pew-Stewart Scholar.

## AUTHOR CONTRIBUTIONS

Conceptualization, K.B. and G.U.; Methodology, G.U. and K.B.; Formal Analysis, G.U.; Investigation, G.U., B.P., R.E., H.-W.Y and E.C.B; Writing – Original Draft, G.U. and K.B.; Funding Acquisition, K.B. and G.U.

## SUPPLEMENTARY FIGURE LEGENDS

**Figure S1. Positive selection CRISPRa screens reveal *SLCO2B1* as the common top hit providing resistance to iron restriction in human and mouse cells**

Related to Figures 1 and 2

(A) Immunoblot analysis for Cas9 expression in PaTu-8988t and Jurkat single cell clones. Clones highlighted in red were used for screens

(B) Top 15 genes scoring as protective against palmitate toxicity in Jurkat CRISPRa screens.

Differential gene scores (palmitate – untreated control) are plotted. Lipid metabolism-related genes are highlighted in orange.

(C) Gene score ranks from BafA1 and DFO CRISPRa screens in Jurkat cells. SLCO2B1 is highlighted as it has the lowest common rank, i.e. highest score.

(D) *SLCO2B1* sgRNA scores in CRISPRa screens for DFO in Jurkat cells

(E) *SLCO2B1* sgRNA scores in CRISPRa screens for BafA1 in Jurkat cells

(F) Immunoblot analysis for SLCO2B1 expression in Jurkat cells expressing vector or *SLCO2B1* cDNA. β-actin was used as loading control.

(G) Fold change in the number (log_2_) of vector, *SLCO2B1* long isoform and *SLCO2B1* short isoform expressing Jurkat cells after 5 days in the presence of indicated BafA1 concentrations.

(H) Fold change in the number (log_2_) of vector, *SLCO2B1* long isoform and *SLCO2B1* short isoform expressing Jurkat cells after 5 days in the presence of indicated DFO concentrations.

(I) Fold change in the number (log_2_) of vector, *SLCO2B1* long isoform and *SLCO2B1* short isoform expressing PaTu-8988t cells after 5 days in the presence of indicated BafA1 concentrations.

(J) Fold change in the number (log_2_) of vector, *SLCO2B1* long isoform and *SLCO2B1* short isoform expressing PaTu-8988t cells after 5 days in the presence of indicated DFO concentrations.

(K) Gene scores of untreated vs. BafA1 (15 nM)-treated mouse AK196 pancreas-dCas9a cells. Gene score is the median log_2_ fold change in the abundance of all sgRNAs targeting that gene during the screening period.

(L) *Slco2b1* sgRNA scores in positive selection CRISPRa screens for BafA1 treatment in mouse AK196 pancreas cells

(M) Gene scores of untreated vs. ConA (1 nM)-treated mouse AK196 pancreas-dCas9a cells.

(N) *Slco2b1* sgRNA scores in positive selection CRISPRa screens for ConA treatment in mouse AK196 pancreas cells

(O) Gene score ranks from BafA1 and ConA CRISPRa screens in mouse AK196 pancreas cells. Slco2b1 is highlighted as the lowest common rank, i.e. highest score.

**Figure S2. *SLCO2B1* overexpression leads to an increase in cellular iron availability**

Related to Figure 2

(A-C) Volcano plots showing the log_2_ fold change in polar metabolite abundance between *SLCO2B1* and *SLCO2A1*-expressing Jurkat cells vs. -log P values. The dotted lines on x-axis indicate 2-fold difference; that on y-axis represents P = 0.05 (Student’s t-test, two-tailed, 95% confidence interval). (A) untreated, (B) DFO (3 μM)-treated and (C) BafA1 (3nM)-treated cells were assayed.

(D) Immunoblot analysis of FTH1 and SLCO2B1 in select human and mouse cell lines expressing vector or *SLCO2B1* cDNA. β-actin was used as loading control.

(E) Immunoblot analysis of TFRC in *TFRC* knockout Jurkat cells expressing vector, *SLCO2A1* or *SLCO2B1* cDNA. β-actin was included as loading control for immunoblots.

**Figure S3. *SLCO2B1* expression mediates resistance to iron restriction through heme oxygenase and overcomes heme deficiency**

Related to Figure 3

Gene score ranks from BafA1 screens in parental (3 nM BafA1) and SLCO2B1-expressing (4nM BafA1) Jurkat cells. Common hits essential in BafA1 treatment, regardless of the genotype, are in yellow highlighted area (genes scoring within top 32 in both screens). Genes that are only essential in *SLCO2B1*-overexpressing cells for BafA1-resistance are in green highlighted area. *HMOX2*, shown in green circle, is one of the top hits specifically in *SLCO2B1*-overexpressing cells.

(B) Top 50 genes scoring as differentially essential upon BafA1 treatment in *SLCO2B1*-expressing cells were grouped in metabolic pathways in which they are functioning.

(C) Fold change in the number (log_2_) of *HMOX2* knockout Jurkat cells expressing vector, *SLCO2A1* or *SLCO2B1* cDNA, during 5-day incubation with or without DFO at indicated concentrations. (Mean ± SEM, n= 3, Student’s t-test, 95% confidence interval).

(D) Schematic depicting key reacting in heme biosynthesis and heme degradation pathways. Succinyl acetone is a potent inhibitor of heme biosynthesis, targeting ALAS1.

(E) *TFRC* sgRNA scores in CRISPR knockout screens of Jurkat cells expressing a control vector or *SLCO2B1* cDNA.

(F) *UROD* sgRNA scores in CRISPR knockout screens of Jurkat cells expressing a control vector or *SLCO2B1* cDNA.

(G) Immunoblot analysis for UROD and ALAS1 in UROD knockout HepG2 cells expressing a control vector or SLCO2B1 cDNA. The position of ALAS1 band was shown with an arrow. β-actin was used as loading control.

(H) Immunoblot analysis of SLCO2B1 and ALAS1 in untreated control and 1 mM succinyl acetone-treated HepG2 cells expressing a control vector or *SLCO2B1* cDNA. β-actin was used as loading control.

**Figure S4. SLCO2B1 localizes to plasma membrane**

Related to Figure 4

(A) Confocal microscopy images of PaTu-8988t cells expressing SLCO2A1-eGFP or SLCO2B1-eGFP. Cell were immunostained for eGFP and LAMP1 (lysosomal marker). DAPI was used as nuclear counterstain. PM: plasma membrane.

(B) Immunoblot analysis for eGFP in PaTu-8988t cells expressing a control vector, SLCO2A1-eGFP or SLCO2B1-eGFP, either untreated or treated with indicated concentration of BafA1. β-actin was used as loading control.

(C) Fold change in the number (log_2_) of PaTu-8988t cells expressing a control vector, SLCO2A1-eGFP or SLCO2B1-eGFP cDNA, during 5-day incubation with or without BafA1 at indicated concentrations. (Mean ± SEM, n= 3, Student’s t-test, 95% confidence interval).

(D) Immunoblot analysis for eGFP and FTH1 in PaTu-8988t cells expressing a control vector, SLCO2A1-eGFP or SLCO2B1-eGFP cDNA. β-actin was used as loading control.

(E) Flow cytometry analysis of ZnMP uptake in mouse HY15549 pancreas cells expressing a control vector or *Slco2b1* cDNA, treated with1 μM ZnMP for 15 min. ZnMP+ gate shown here was applied in dose-dependent ZnMP uptake assays

(F) Relative expression levels of *Slco2b1* and *Slco2a1* genes in wild-type (+/+), heterozygous (+/−) and homozygous mutant (+/−) mice from *Slco2b1^tm1a^* mouse line. Expression levels were normalized to beta-actin. Specific loss of *Sloc2b1* expression in homozygous mutants was observed; *Slco2a1* expression was assayed as a control. (Mean ± SEM, n= 3, Student’s t-test, 95% confidence interval).

**Table 1. Reagents and resources**

**SUPPLEMENTARY TABLES**

Table S1. Antimycin A CRISPRa screen results in PaTu-8988t cells

Table S2. CB-839 CRISPRa screen results in PaTu-8988t cells

Table S3. BSO CRISPRa screen results in PaTu-8988t cells

Table S4. Palmitate CRISPRa screen results in Jurkat cells

Table S5. Differential Gene Scores in Palmitate CRISPRa screen performed in Jurkat cells

Table S6. Gene Scores from Jurkat CRISPRa screens for DFO

Table S7. Differential Gene Scores in DFO CRISPRa screen performed in Jurkat cells

Table S8. Gene Scores from Jurkat CRISPRa screens for BafA1

Table S9. Differential Gene Scores in BafA1 CRISPRa screen performed in Jurkat cells

Table S10. Gene Scores from CRISPRa screens for BafA1 in mouse AK196 pancreas cells

Table S11. Gene Scores from CRISPRa screens for ConA in mouse AK196 pancreas cells

Table S12. Gene Scores from CRISPR knockout screens for BafA1 in *SLCO2B1*-expressing Jurkat cells

Table S13. Differential Gene Scores from CRISPR knockout screens for BafA1 in *SLCO2B1*-expressing Jurkat cells

Table S14. Gene Scores from CRISPR knockout screens for BafA1 in parental Jurkat cells

Table S15. Gene Scores from CRISPR knockout screens for differential essentiality of genes between control vector and *SLCO2B1*-expressing Jurkat cells

