## Supplementary figures and images for "Metabolic-scale gene activation screens identify SLCO2B1 as a heme transporter that enhances cellular iron availability"

### Figure S1

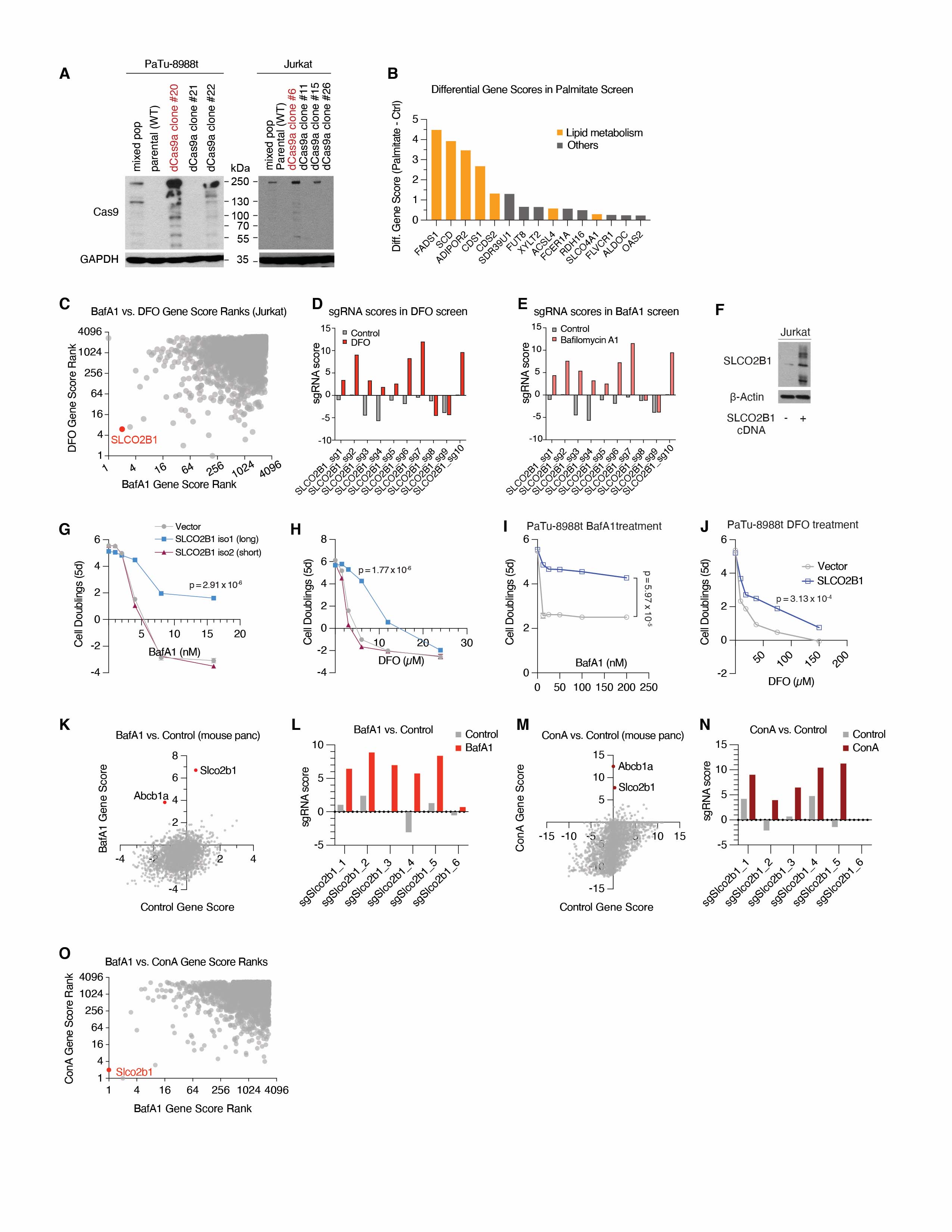

### Figure S2

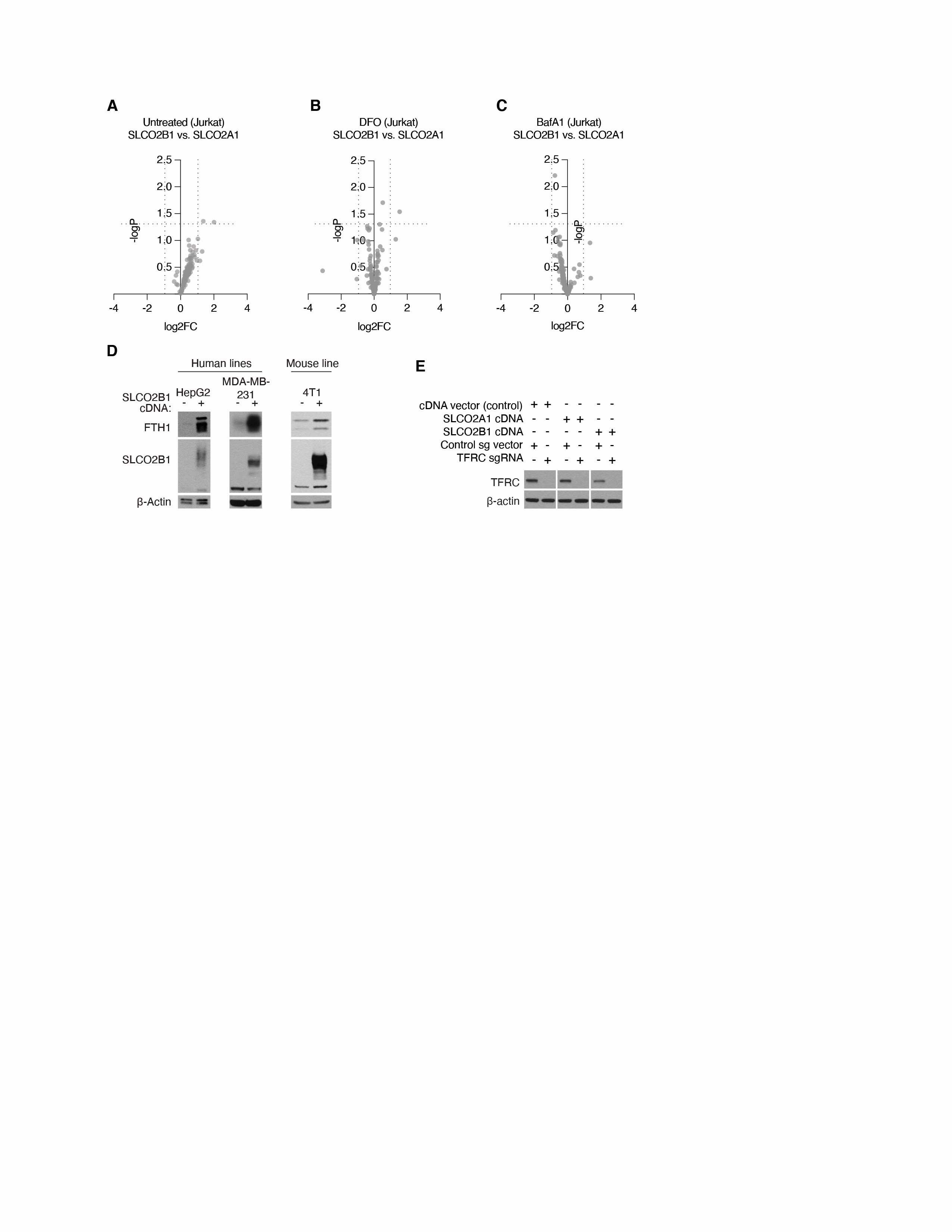

### Figure S3

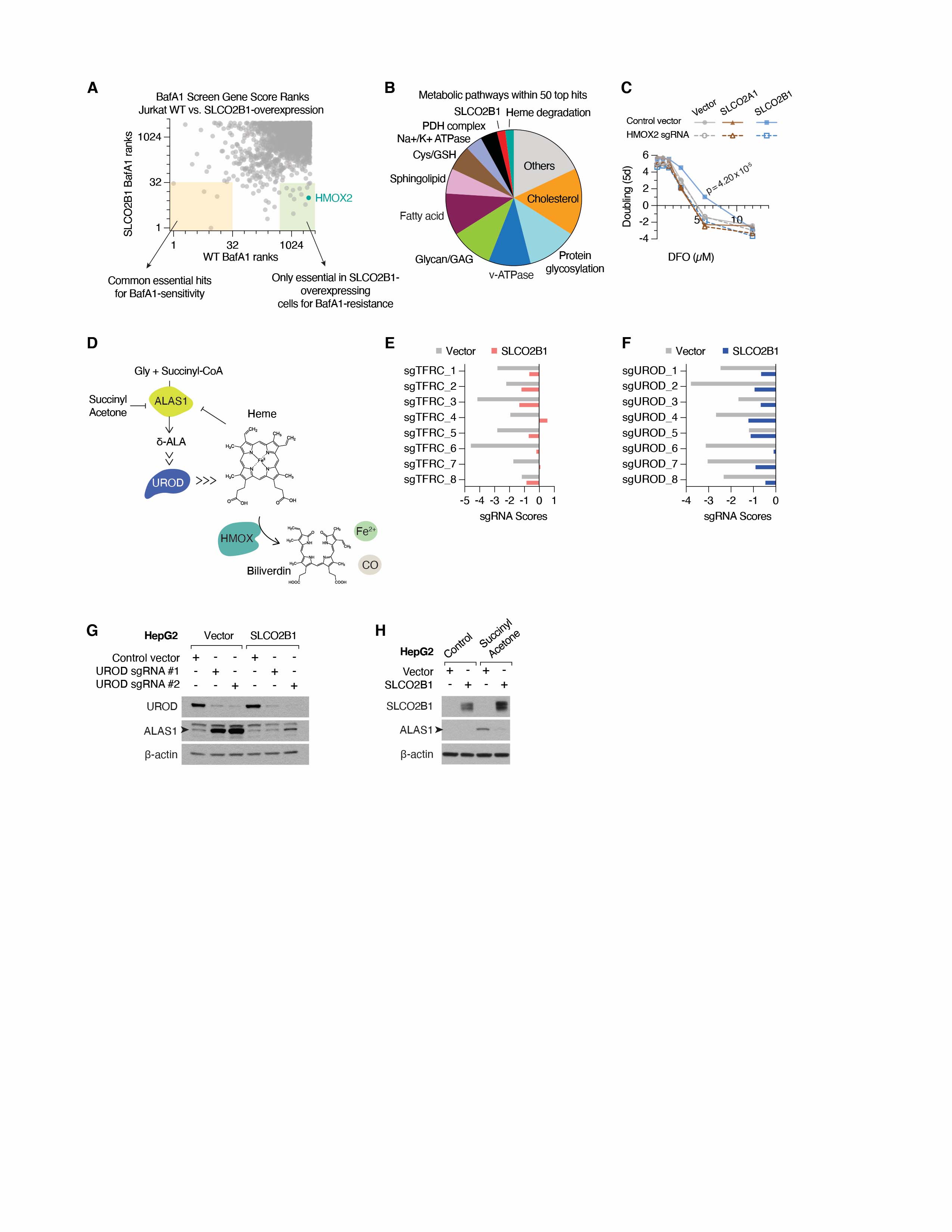

### Figure S4

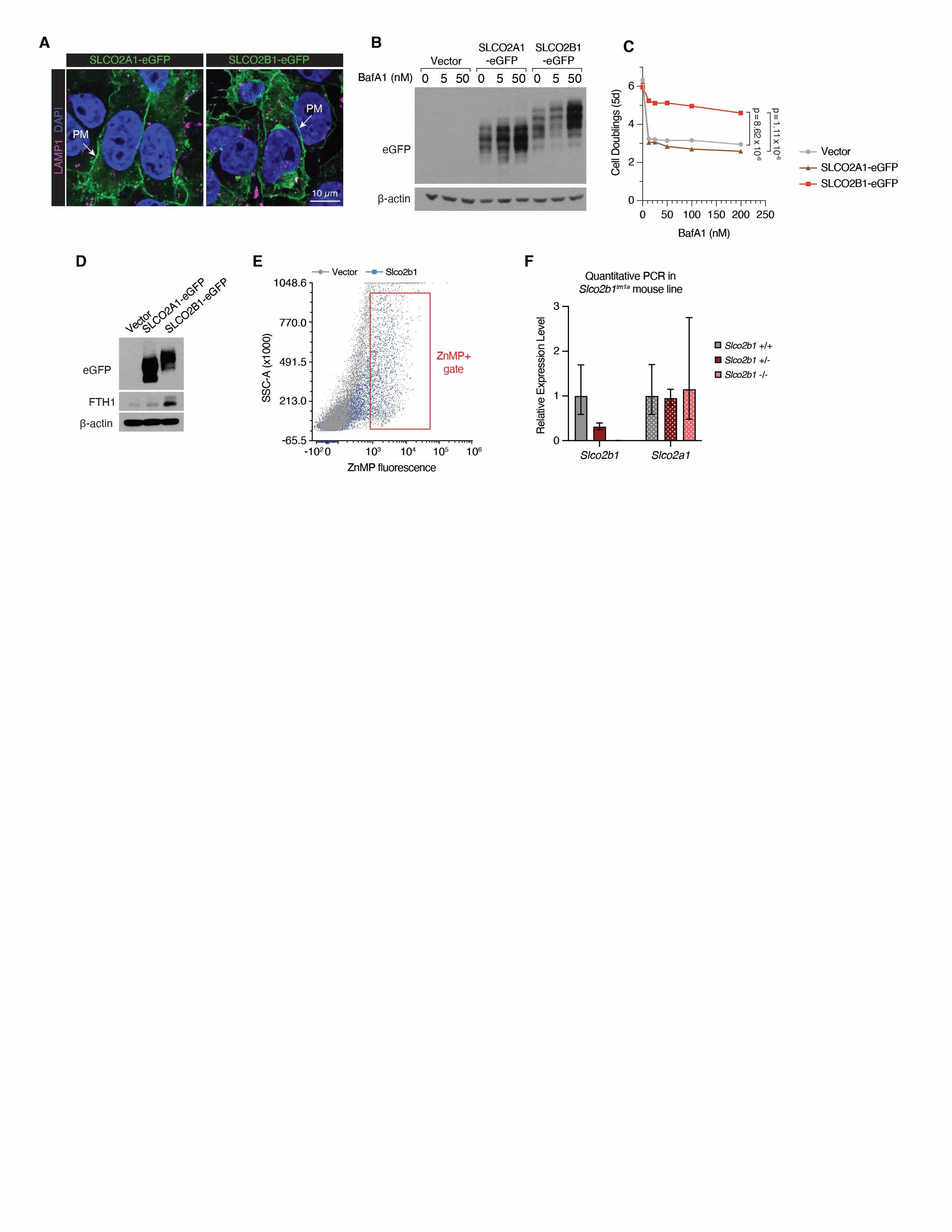
